# Predicting cross-tissue hormone-gene relations using balanced word embeddings

**DOI:** 10.1101/2021.01.28.428707

**Authors:** Aditya Jadhav, Tarun Kumar, Mohit Raghavendra, Tamizhini Loganathan, Manikandan Narayanan

## Abstract

**Motivation:** Large volumes of biomedical literature present an opportunity to build whole-body human models comprising both within-tissue and across-tissue interactions among genes. Current studies have mostly focused on identifying within-tissue or tissue-agnostic associations, with a heavy emphasis on associations among disease, genes and drugs. Literature mining studies that extract relations pertaining to inter-tissue communication, such as between genes and hormones, are solely missing.

**Results:** We present here a first study to identify from literature the genes involved in inter-tissue signaling via a hormone in the human body. Our models BioEmbedS and BioEmbedS-TS respectively predict if a hormone-gene pair is associated or not, and whether an associated gene is involved in the hormone’s production or response. Our models are classifiers trained on word embeddings that we had carefully balanced across different strata of the training data such as across production vs. response genes of a hormone (or) well-studied vs. poorly-represented hormones in the literature. Model training and evaluation are enabled by a unified dataset called HGv1 of ground-truth associations between genes and known endocrine hormones that we had compiled. Our models not only recapitulate known gene mediators of tissue-tissue signaling (e.g., at average 70.4% accuracy for BioEmbedS), but also predicts novel genes involved in inter-tissue communication in humans. Furthermore, the species-agnostic nature of our ground-truth HGv1 data and our predictive modeling approach, demonstrated concretely using human data and generalized to mouse, hold much promise for future work on elucidating inter-tissue signaling in other multi-cellular organisms.

**Availability:** Proposed HGv1 dataset along with our models’ predictions, and the associated code to reproduce this work are available respectively at https://cross-tissue-signaling.herokuapp.com/, and https://github.com/BIRDSgroup/BioEmbedS.

## 1 Introduction

Inter-tissue communication forms the basis for life and health in multi-cellular organisms, and complex diseases often affect multiple organs/tissues. A grand goal of systems biology is to develop whole-body models that capture not only within-tissue biomolecular interactions but also across-tissue interactions. Recently developed whole-body *metabolic* models for humans [5] (such as the Harvey and Harvetta models encompassing 26+ organs and their metabolic interactions culled from literature, and refined by large-scale genomic or other experimental data [25]) are promising, but models of similar scale at the *gene-level* are lacking. Emerging multi-tissue genomic datasets (like the NIH GTEx data [10]) have encouraged some efforts to address this gap, and there is an urgent need to augment/validate such data-driven whole-body gene networks by literature-driven approaches. An ideal literature mining system would extract relations of the form “Gene X in Tissue A interacts with Gene Y in Tissue B via a mediating signaling molecule”.

In this work, we harness the large volume of biomedical literature (present in repositories like PubMed [27] with ∼23 million abstracts) to identify genes involved in inter-tissue signaling mediated by hormones. We specifically focus on endocrine hormones, as they are a popular class of signaling molecules and endocrine biology has revealed and continues to reveal many hormones and their regulating genes. For example, in the pancreas tissue, gene *INS* produces the well-studied insulin hormone, which is processed and secreted into the blood with the help of other gene products and biomolecules; and the protein encoded by the *INSR* gene is the primary receptor of the hormone in muscle and other tissues, where *INSR* gets activated by the insulin hormone and affects several other genes downstream to regulate glucose uptake by body cells. A more recent example is a new type of vesicle-mediated communication between liver and adipose tissues involving non-protein-coding (microRNA) genes [8, 32]. The goal of this work is to extract all such hormone-gene relations, and classifying the hormone-associated genes as either source (genes aiding in hormone production/processing/secretion at a source tissue) or target (genes responding to the hormone directly via binding or as a downstream response at a target tissue).

Several challenges stand in the way of our goal of extracting hormone-gene relations from literature, including:

1. Lack of a unified database of ground-truth hormone-gene associations, since current hormone databases like HMRbase [23] and EndoNet [11] focus on the primary gene coding for a (peptide) hormone and the primary receptor genes, and not on the many other genes involved in hormonal processing/response.
2. Severe imbalance in known hormone-gene relations both in the number of source vs. target genes of each hormone, and of associated genes per hormone which varies widely across hormones (e.g., insulin is better studied than many other hormones).
3. Lack of standard *in silico* strategies for large-scale validation of novel hormone-gene predictions using independent data on inter-tissue signaling.

Our work attempts to address the barriers above to characterize inter-tissue signaling, and current studies have addressed only the second challenge above and that too in very different contexts/applications. Specifically, current literature mining studies largely extracted relations among entities that are tissue-agnostic (i.e., doesn’t depend on the tissue) or within-tissue (i.e., happens within one or more of the tissues or cell types), and includes relations such as disease-gene [1], gene-phenotype [28], drug-drug [29], or protein-protein/gene-gene [30, 24] types, or subsets of them [4, 15]. Besides using traditional NLP (Natural Language Processing) features based on syntax/semantics, these methods also exploit modern advances like word embeddings, which are vector representations of words that are learnt via deep learning models (Word2Vec [20] or FastText [2]) to capture the semantic similarities and relationships among words. Examples include a joint ensemble learning approach by Bhasuran et al. [1] for disease-gene predictions, which builds upon a Support Vector Machine (SVM) classifier called BeFree introduced earlier by Bravo et al. [4]; and an approach by Park et al. [21] that combines traditional NLP techniques with Word2Vec embeddings.

Given this context, our work on systematic prediction of genes mediating inter-tissue signaling makes three main contributions:

1. Our work is the first study to predict hormone-gene associations from biomedical literature, with our focus on inter-tissue communication setting it apart from earlier literature mining studies on predicting tissue-agnostic or within-tissue interactions.
2. Our work is enabled by expressly compiling a ground-truth bipartite (hormone-gene) database HGv1, and balancing it in the space of mapped word embeddings to avoid well-studied hormones and source-vs-target genes’ imbalance from unduly influencing our model predictions.
3. Our models BioEmbedS and BioEmbedS-TS, which are SVM classifiers trained on these balanced word embeddings, not only corroborates existing hormone-gene links and hormone source vs. target genes (collated in HGv1, respectively at an average accuracy of 70.4% and 72.5%), but also predicts novel gene associations of hormones. These novel genes are enriched for diseases known to be related to the corresponding hormone, across many different hormones.

## 2 Methods

### 2.1 Assembling a ground-truth dataset: HGv1

Since a unified database of source and target genes for known hormones was not available, we expressly assembled such a database for 51 endocrine hormones, primarily ones listed in a Endocrine Society website^1^, by integrating data from several sources [9, 23, 11]. We manually went through all GO (Gene Ontology [9]) term names mentioning a given hormone, and manually extracted the GO terms that could be unambiguously added to source and target sets for the hormone (see Table 1). Every GO term we considered represents a species-agnostic biological process or molecular function, and can hence be tailored to any species by taking the appropriate set of genes annotated to the term. We compiled a human dataset HGv1.human of about two thousand hormone-gene associations (Table 2) derived from the appropriate GO terms and augmented with primary genes (genes encoding a peptide hormone or hormone-binding receptors) from other sources [23, 11]. Similarly, we constructed HGv1.mouse dataset by collecting mouse genes annotated to the GO terms collected in the first phase, and by taking the mouse homologs^2^ of the primary human genes. This work focuses on HGv1.human (simply referred to as HGv1 in the text), with HGv1.mouse dataset being used to study generalizability of our models.

**Table 1:** Example snapshot of our HGv1 dataset: Glucagon and Adiponectin hormones along with their source and target GO terms.

| Hormone | Source terms | Target terms |
| --- | --- | --- |
| Glucagon | GO:0070091<br>(Glucagon secretion) | GO:0033762 (response to Glucagon) |
|  | GO:0120116<br>(Glucagon processing) | GO:0004967 (Glucagon receptor activity) |
|  |  | GO:0031769 (Glucagon receptor binding) |
| Adiponectin | GO:0070162<br>(Adiponectin secretion) | GO:0055100 (Adiponectin binding) |
|  |  | GO:0033211 (Adiponectin-activated signaling pathway) |

**Table 2:** HGv1 dataset overview: Summary of the 2, 009 hormone-gene associations in our HGv1.human dataset. Note ± denotes standard deviation here and elsewhere in the text.

|  |  |
| --- | --- |
| Number (#) of hormones/genes | 51 / 1453 |
| Median # of genes associated with a hormone | 39.39 (± 78.36) |
| Median # of hormones associated with a gene | 1.38 (± 0.86) |
| # of source/target genes across all hormones | 519 / 1082 |
| Median # of source genes for a hormone | 14.3 (± 33.46) |
| Median # of target genes for a hormone | 28.8 (± 54.04) |

<sup>1</sup>https://www.hormone.org/your-health-and-hormones/glands-and-hormones-a-to-z, accessed Jul 23, 2019.
<sup>2</sup>Human-mouse homology mapping done via MGI Batch Query (http://www.informatics.jax.org/batch), accessed Nov 11, 2020.

### 2.2 Our BioEmbedS and BioEmbedS-TS approach

#### Word embeddings based classifiers

We develop two classification models – BioEmbedS to predict hormone-gene associations, and BioEmbedS-TS to classify an associated gene into source vs. target set of a hormone. These models, referred jointly as BioEmbedS*, are trained and evaluated using the HGv1 dataset, and use word embeddings as input features (Figure 1). Specifically, we use word embeddings for gene symbols and hormones from a FastText model pretrained on the PubMed biomedical corpus called BioWordVec^3^ [31]. FastText [2] is a neural network model learned by minimizing cross-entropy loss between each word and its predicted context within a fixed window size. We used 200-dimensional embeddings obtained using a skip-gram based implementation with window size 20. This ensures the embedding vectors of not only co-occurring entities in a document but also entities with similar word neighborhoods exhibit high similarity [21]. BioWordVec actually uses subword information to obtain embeddings of words (including out-of-vocabulary words), i.e., the embedding of each word is represented by the sum of embeddings of all n-grams (3≤ n≤ 6) in the word (after converting the words, including hormones and gene symbols, to lower-case) [31, 2]. Please note that BioWordVec is already trained using the PubMed corpus and so our HGv1 dataset is not used to obtain the word embeddings; HGv1 (training/testing splits) is instead used along with the pre-trained word embeddings to build our BioEmbedS* classifiers.

**Figure 1:**
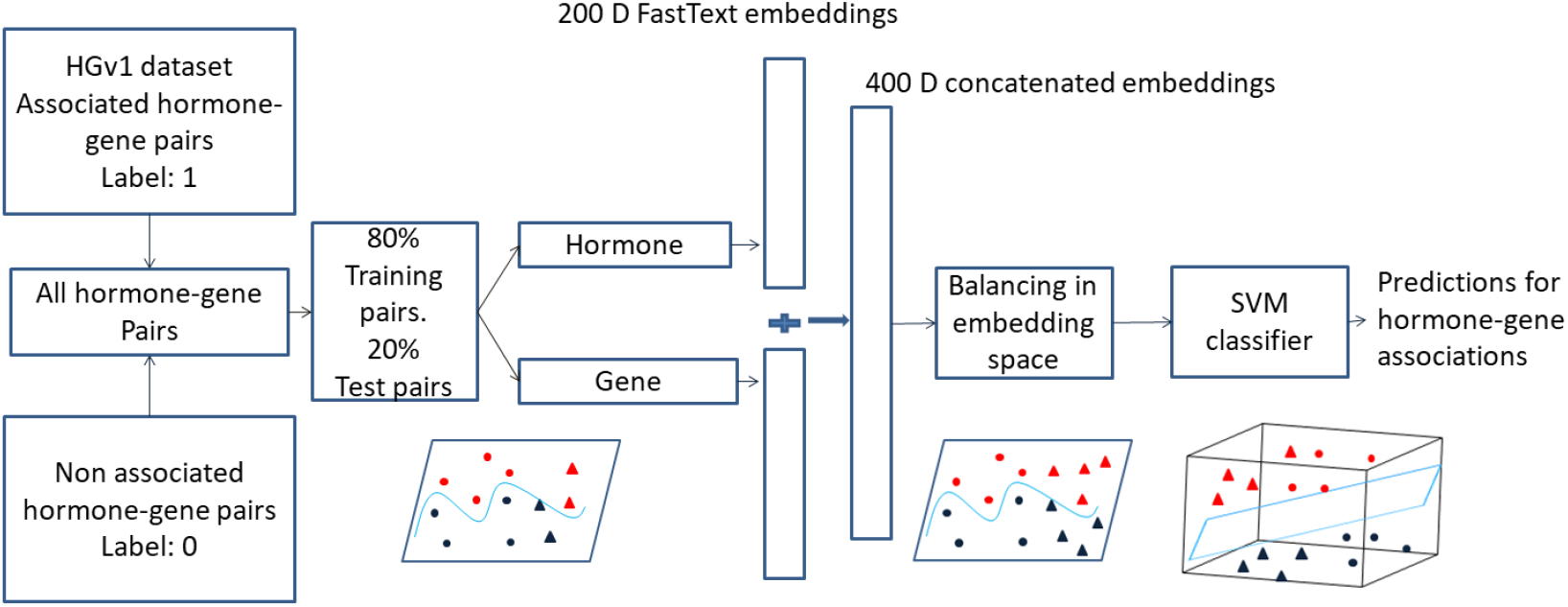
BioEmbedS model overview: Our BioEmbedS model predicts if a hormone-gene pair is associated or not from D-dimensional word embedding vectors of the hormone name and the gene symbol. Our HGv1 dataset is crucial for proper training/evaluation of our model. In the toy-example shown, circles and triangles indicate hormone-gene pairs for two illustrative hormones; and blue and red colors respectively denote the positive (associated) and negative (non-associated) genes for each hormone. The positive/negative classes are balanced across the two hormones, before separating them in a higher dimensional space using a SVM classifier. BioEmbedS-TS model has source and target genes for a hormone in place of positive and negative genes.

Existing studies on prediction of disease-gene associations have shown that SVM (Support Vector Machines) classifiers perform well when used with word embeddings [1]. So we explored SVM along with RF (Random Forest) classifiers as our primary classifiers, and compared them with other secondary choices of classifiers as well, and found SVMs to perform well (see Results). Hence we decided on a SVM based model to predict hormone-gene relations and call it BioEmbedS (with S denoting SVM). Similarly, we use the SVM based model to classify source vs. target genes for associated genes for a given hormone and call it BioEmbedS-TS (with - TS denoting Target vs. Source).

#### Stratified/nested CV (cross-validation)

We explored the parameter space (both hyper-parameter tuning and parameter learning) of our SVM and RF models using a 5-fold CV strategy that is stratified to ensure even distribution of each hormone’s genes across the different folds, and nested to allow proper partitioning of the HGv1 data into training, validation and test sets. In detail, a stratified split amounts to considering each hormone with a certain number of associated genes in HGv1 (denoted *n*), and putting randomly chosen 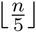 genes into each of the 5 folds, and the remainder genes randomly (one each) into any of the 5 folds. This procedure done for the BioEmbedS model ensures that each fold has genes belonging to each hormone proportional to their presence in the overall dataset. A similar procedure to distribute the number of source (and separately target) genes of each hormone evenly across the 5 folds was done for the BioEmbedS-TS model also. In the nested 5-fold CV strategy that we use to build both the models, we make a train/validation/test split using 3/1/1 folds respectively, and identify the classifier’s parameters by using only train and validation splits. The test split is kept aside from the training process, and used solely to report the model performance.

### 2.3 Balancing word embeddings within the nested CV framework

#### Over/under-sampling (balancing) training set embeddings

Our HGv1 dataset has a skewed distribution of the number of genes associated with different hormones (Table 2); so it is important to prevent well-studied hormones (with a large number of gene associations in HGv1) from unduly influencing our model. To address the problem of class imbalance, oversampling techniques like SMOTE (Synthetic Minority Oversampling Technique) [6] synthesize new examples from existing ones for the minority class, whereas undersampling techniques like Condensed Nearest Neighbours [14], TOMEK Links [26], etc. selectively remove examples from the majority class. A combination of oversampling and undersampling techniques is shown to perform better than using either alone [6]. We strategically apply a combination of SMOTE and TOMEK Links on the mapped embeddings of only genes with exactly one hormone association in HGv1. Working in the space of embeddings of such “uni-hormone” genes is both desirable in facilitating SMOTE oversampling of our data, and permissible as a large fraction of all genes in HGv1, 1102 of all 1453, are uni-hormone (i.e., associated uniquely with some hormone, as opposed to being multi-hormone or associated with more than one hormone within HGv1). This technique applied to BioEmbedS and a similar technique applied to BioEmbedS-TS handle the large variation in HGv1 in the number and type (source vs. target) of genes respectively across hormones.

For BioEmbedS, we assign a unique class ID to every hormone, and all the genes uniquely associated with the hormone belong to the class indicated by its class ID (with all multi-hormone genes discarded). We then use a combination of SMOTE oversampling and TOMEK Links undersampling (with the number *k* of nearest neighbors in SMOTE set to 2 in the implementation used [18]) to get an approximately equal number of genes/examples (same as the highest number of genes associated with a hormone among all the hormones in HGv1) for every hormone. We now have a dataset with approximately equal numbers of the genes related to every hormone that forms hormone-gene pairs of positive class for our binary classification problem. The following strategy is applied to create the negative class (set of non associated hormone-gene pairs). For every hormone, we construct a set that contains the genes (synthesized examples from oversampling and undersampling) associated with all the hormones except the one under consideration. To maintain the class balance between positive and negative associations for a hormone, we select as many examples present for the positive set of that hormone, from this set. Repeating this process for each hormone results in a set of negative hormone-gene associations for each hormone. Finally, we have a balanced dataset across different hormones with positive/negative classes.

For BioEmbedS-TS, we define two unique IDs for every hormone; one maps to the source genes and the other maps to the target genes for that hormone (again we focus only on genes with exactly one hormone association, and additionally remove the small set of genes that are annotated as both source and target for the same hormone). The source and target genes associated with a hormone belong to classes indicated by their IDs. This makes the number of classes equal to twice the number of hormones present in our dataset. We use SMOTE and TOMEK Links strategy as for BioEmbedS to get an approximately equal number of source and target genes (same as the highest number of source/target genes associated to a hormone among all the hormones in HGv1) for every hormone. Finally, we have a dataset that is balanced across the different hormones and source/target class.

#### Balancing and model selection within the nested CV framework

The overall careful application of our balancing and model selection steps to the 3/1/1 train/validation/test folds is shown in Algorithm 1, along with what constitutes the training and testing sets of each inner/outer CV iteration (e.g., validation and test folds are the respective testing sets for inner and outer CV loops). As for the genes considered by Algorithm 1, training sets are restricted to contain only uni-hormone genes to facilitate application of SMOTE and TOMEK Links as discussed before, whereas testing sets are allowed to contain both uni- and multi- hormone genes to reflect a real-world setting where any gene may be queried for its association to a hormone. As for the hormones considered, the overall algorithm considers only hormones with at least 5 gene associations to permit 5-fold CV; additionally, each inner/outer CV iteration considers only hormones with sufficient gene associations as “eligible” for further analysis. In detail, every hormone with at least 3 associations to uni-hormone genes in an iteration’s training set is considered eligible for this iteration (to permit 2-nearest-neighbor SMOTE), and the remaining hormones are removed from this iteration’s training as well as testing sets.

SVM and RF models with different hyper-parameter combinations are trained on the resulting balanced training folds, and Cohen’s Kappa score of these trained models on the validation folds is used to select the best model. In each iteration, the test fold is never used for training or choosing hyper-parameters for the classifier. For BioEmbedS, a SVM model with a third degree polynomial kernel was chosen as the best classifier by Algorithm 1 (Step 10) consistently in all 5 outer CV iterations. This SVM model was later trained using the entire HGv1 dataset, after applying similar restrictions and balancing as in Step 11 of Algorithm 1, to obtain the final BioEmbedS model – this final model was used for making all protein-coding gene predictions (including novel ones discussed in Results). Hormones found eligible for building the final model amounted to 34 and are known as primary hormones hereafter; the remaining 17 ineligible hormones would also have been ineligible (and hence discarded or unseen when choosing/training models) in each of the inner/outer CV iterations of Algorithm 1 due to these hormones’ insufficient gene associations.

**Algorithm 1:**
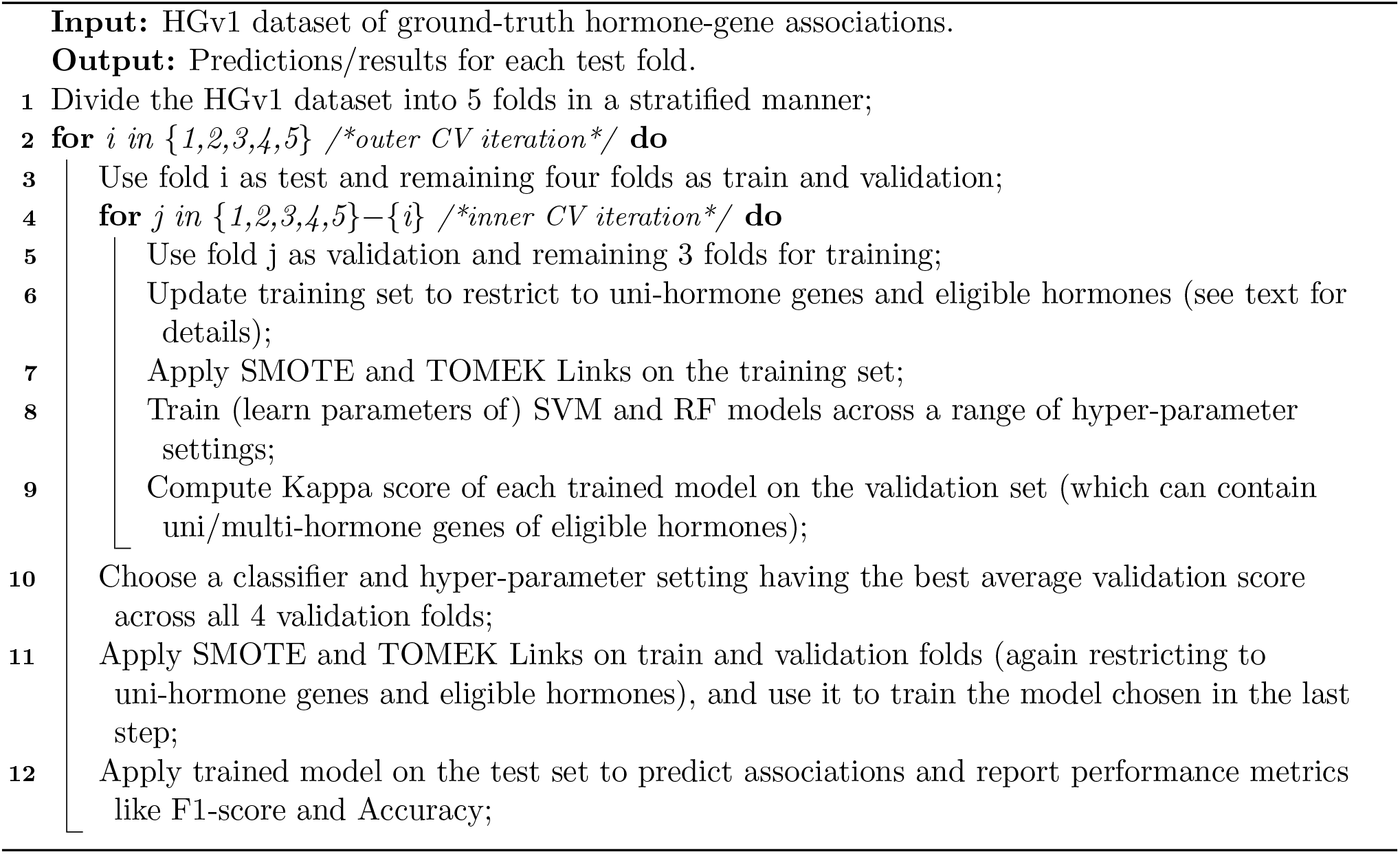
Pseudocode for nested 5-fold CV (cross validation).

### 2.4 Performance metrics, code availability, and disease enrichment analysis

We evaluate our model on five different unseen test sets (as shown in Algorithm 1), by reporting the performance of classifiers on these test sets using standard performance metrics like Precision, Recall, F1-score, Accuracy, Kappa score, Area under the Receiver Operating Characteristics curve (ROC-AUC) and Area under the Precision-Recall curve (PR-AUC). All these metrics take values in the range 0 to 1 and hence can be expressed as percentages (with the exception of Kappa score that can take negative values), with higher values indicating better performance. Suppl Methods 1.1 provides definitions for these metrics assuming the two classes in our binary classification problems are as follows: for BioEmbedS, we naturally let positive and negative class to refer respectively to association and non-association of a hormone-gene pair; for BioEmbedS-TS that starts with an associated hormone-gene pair, we arbitrarily let positive and negative class refer to hormone-source gene pair and hormone-target gene pair association respectively. Given these class definitions, a false positive for instance, for BioEmbedS would be a non-associated hormone-gene pair that is wrongly predicted as associated by our method; and for BioEmbedS-TS would be a hormone-target gene pair that is wrongly predicted as hormone-source gene pair by our method.

Implementations are done using Python *Scikit-learn* framework using *decision function()* and *predict proba()* methods of SVC (Support Vector Classification) in *sklearn* respectively to obtain the SVM model scores for all hormone-gene pairs and the probability of association between hormone-gene pairs (probability score of SVM). For reproducibility purposes, we provide hyperparameter choices in Suppl Methods 1.2 and open-source code in a public repository^4^.

Hormone-gene predictions are called using the final BioEmbedS model described above at a SVM probability score of at least 0.7, with this default cutoff value chosen so as to get a reasonable number of predictions for each hormone. To validate the predicted genes of a hormone, we perform a disease enrichment analysis using the Enrichr tool [7] and DisGeNET [22] collection of known disease-related genes. All reported disease enrichment P-values from this analysis are corrected for multiple testing of different DisGeNET disease terms.

## 3 Results

### 3.1 Word embeddings are informative of relationship among hormones and genes

We first assess the quality of literature-based BioWordVec [31] embeddings of hormone names and gene symbols using simple unsupervised learning methods. Hierarchical clustering of the word embeddings of the 51 hormones in our HGv1 dataset (Figure 2) revealed that functionally similar hormones often group together into clusters – for instance, neurotransmitter hormones like serotonin and dopamine are clustered together; and so are steroid hormones with sexual and reproductive functions such as testosterone, estradiol and progesterone.

**Figure 2:**
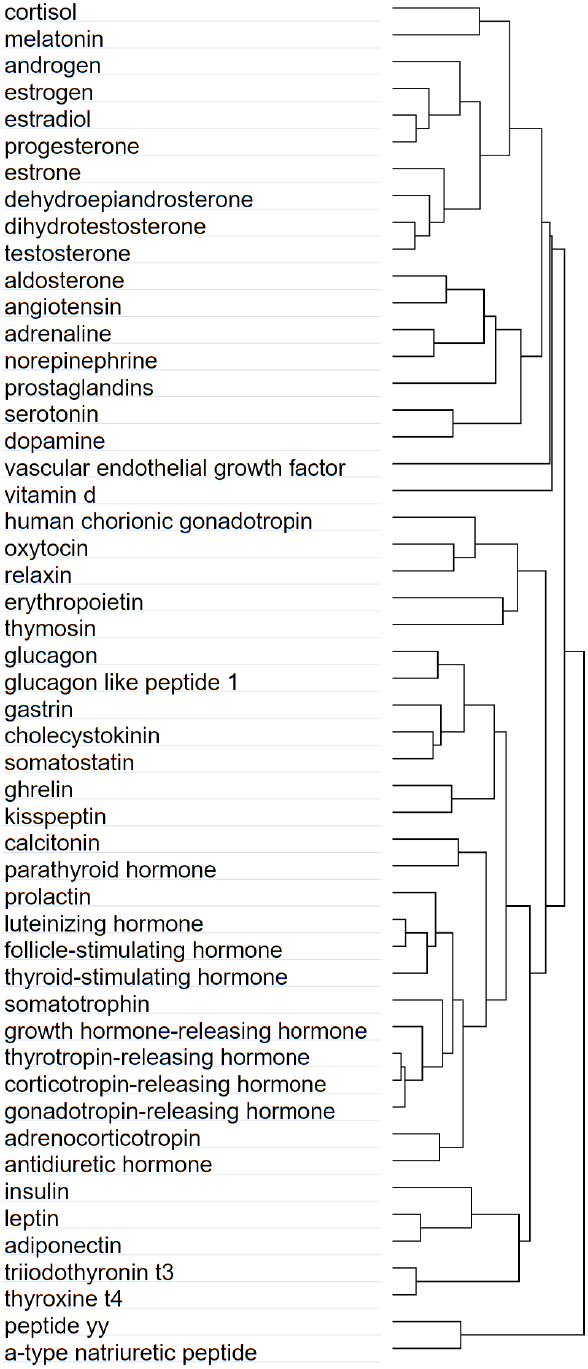
Similarity of hormone embeddings: Hierarchical clustering dendrogram of the 200-dimensional hormone embeddings using complete linkage method and one minus cosine similarity (cosine of the angle between two vectors) as the distance measure.

To evaluate the quality of embeddings of both hormones and genes, we predicted hormone-gene associations using the popular cosine similarity measure between word embeddings of a hormone and a gene, and obtained an average ROC-AUC of 0.69 (Suppl Figure S1). This result confirms the good quality of BioWordVec embeddings, seen in earlier studies on extracting relations among proteins and drugs [31], in our new context of extracting hormone-gene relations from biomedical literature. This result also provides a baseline performance from a simple unsupervised method to compare our supervised BioEmbedS model against.

### 3.2 BioEmbedS strategy on disease-gene predictions is competitive with other methods

Due to lack of existing tools for hormone-gene predictions and due to several tools available for disease-gene predictions, we first validated our BioEmbedS strategy (of a SVM classifier trained on word embeddings) on predicting disease-gene associations from a corpus called EU-ADR [12]. For performance comparison of our method, we used results reported by the BeFree [4] and Joint Ensemble [1] methods discussed before in Introduction. BioEmbedS approach is able to obtain comparable or slightly better F1-score of 85.84% relative to these methods (Table 3). This is promising as our approach, originally conceived for predicting hormone-gene links, performs reasonably well for a disease-gene prediction task. We do not delve further into these specific disease-gene predictions, as this work’s main focus is to predict hormone-gene relations mediating inter-tissue signaling.

**Table 3:**
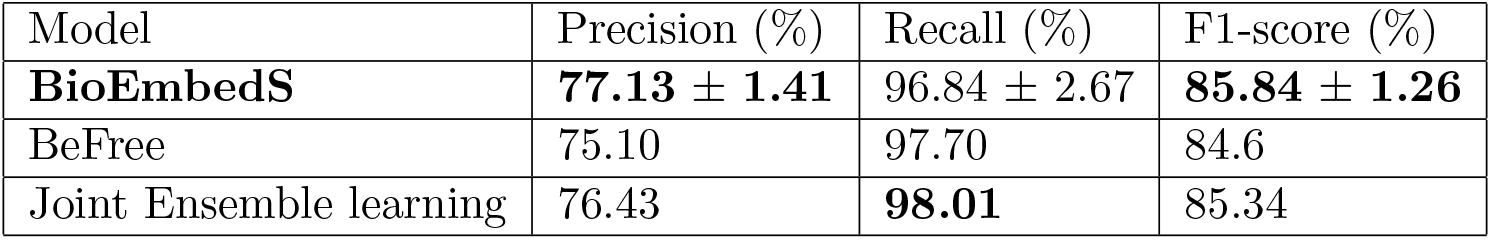
Performance on disease-gene predictions: Our BioEmbedS approach’s 10-fold CV based result is compared against existing methods’ reported results on the EU-ADR disease-gene corpus.

| Model | Precision (%) | Recall (%) | F1-score (%) |
| --- | --- | --- | --- |
| **BioEmbedS** | **77.13 ± 1.41** | 96.84 ± 2.67 | **85.84 ± 1.26** |
| BeFree | 75.10 | 97.70 | 84.6 |
| Joint Ensemble learning | 76.43 | **98.01** | 85.34 |

### 3.3 BioEmbedS predicts hormone-gene pairs reli ably, and better than a generic resource STRING

In our 5-fold CV framework in Algorithm 1, a SVM model with a third degree polynomial kernel was chosen as a consistent classifier, and the resulting BioEmbedS models predicted hormone-gene associations with a reasonably good accuracy of 70.4% ± 1.8% and F1-score of 71.4% ± 2.7% (Suppl Table S1). The SVM based model also achieved better or comparable results than other classifier choices such as logistic regression and decision trees (Suppl Table S2). We also note that in our baseline comparison, our supervised BioEmbedS performed better than the unsupervised cosine similarity approach seen above (Suppl Figure S1).

Since there are no direct hormone-gene prediction tools available to which we can compare our method, the closest alternative was to match a (peptide) hormone to its primary gene encoding the hormone, and then use predicted associations of this gene to other genes. Predicted gene-gene associations are available in a widely-used resource called STRING [24], and we found that our BioEmbedS’ hormone-gene scores were consistently better than STRING’s literature mining based scores for the corresponding (mapped) gene-gene pairs (Figure 3(a)).

**Figure 3:**
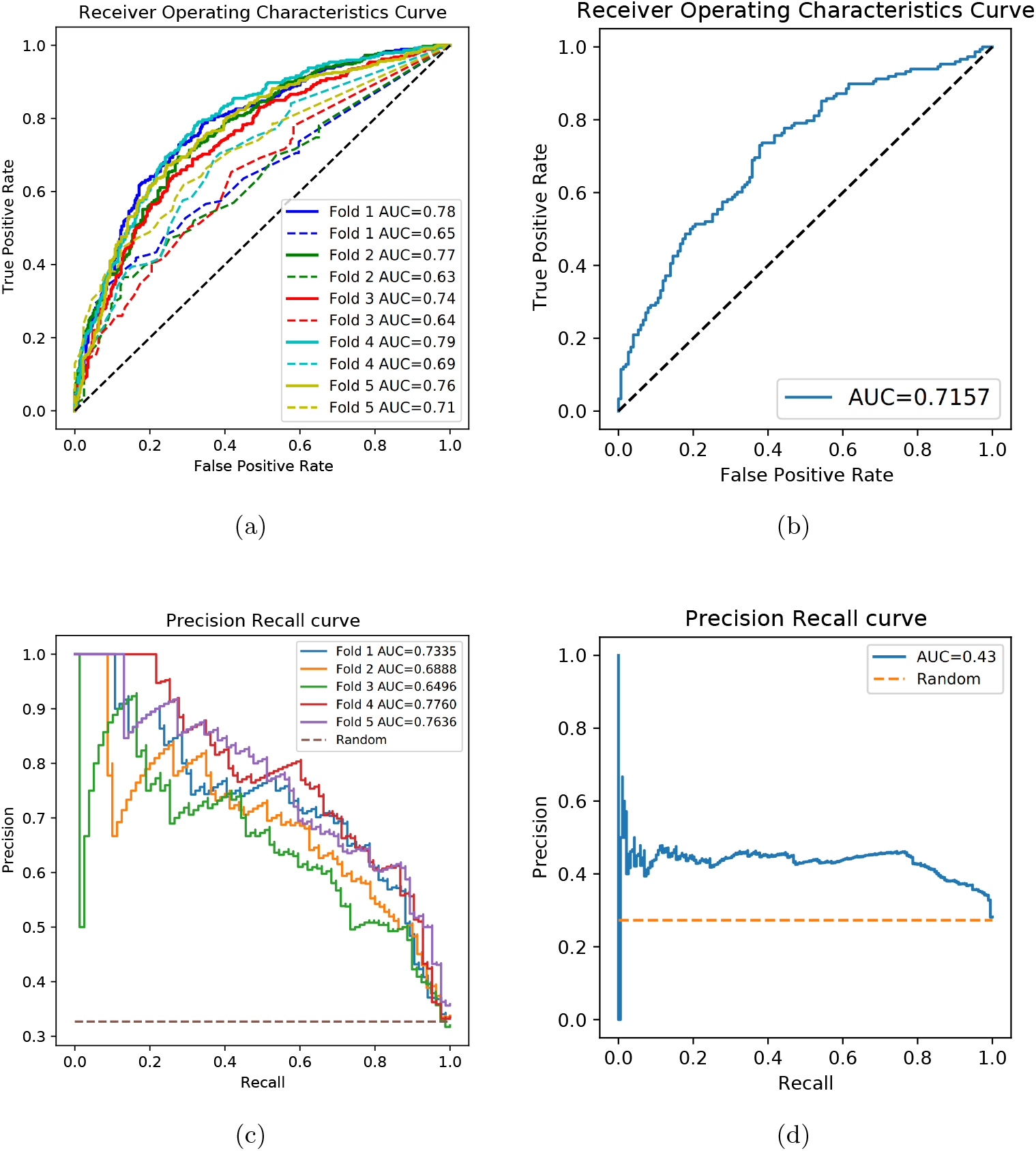
Performance curves of our BioEmbedS* models: (a) ROC curves of BioEmbedS (solid lines) and STRING (dashed lines) for hormone-gene predictions based on 5-fold CV. (b) ROC curve of BioEmbedS for unseen external hormones’ predictions. (c) PR curves of BioEmbedS-TS for source/target gene predictions based on 5-fold CV. (d) PR curve of BioEmbedS-TS for unseen external hormones’ predictions. AUC (Area Under Curve) of a perfect classifier is 1, and a random classifier is 0.5 for ROC curves and specified as a dashed line in the PR curves.

### 3.4 BioEmbedS-TS classifies source vs. target genes across different classes of hormones

For genes known to be associated with a hormone, we next classify it they are source or target genes for the hormone. The performance of our BioEmbedS-TS model was also reasonably good with accuracy of 79% ± 1.9%, and F1-score of 84.8 ± 1.3% for target genes and 66 ± 3.5% for source genes (see Table 4 and Figure 3(c)).

**Table 4:**
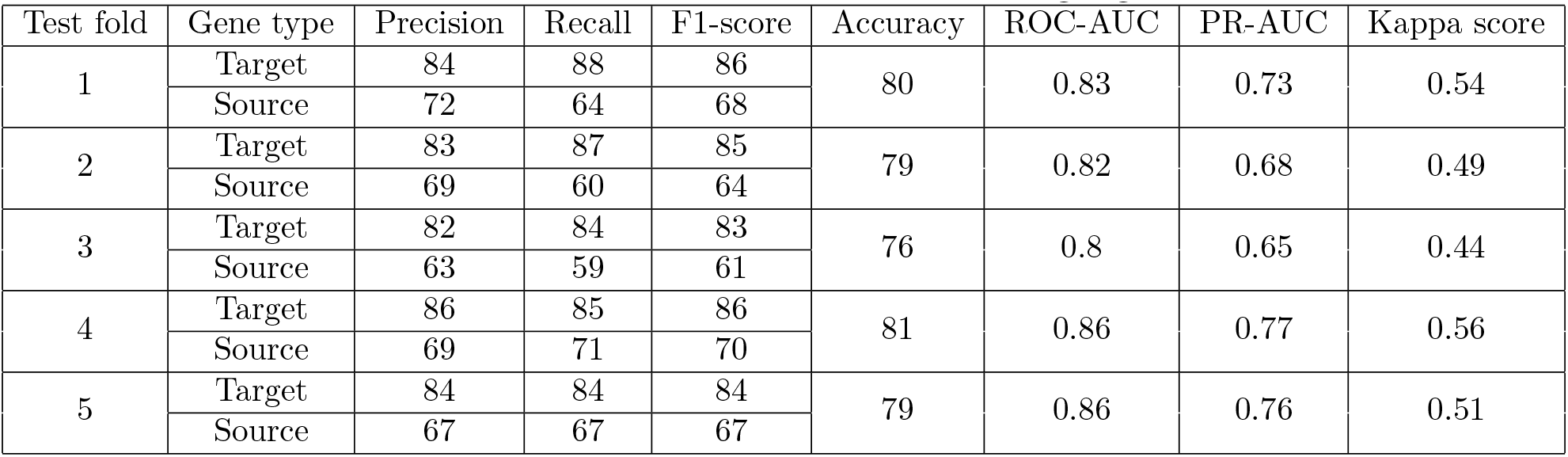
BioEmbedS-TS results: Classification of source vs. target genes across the 5 test sets.

### 3.5 BioEmbedS* models generalize to unseen hormones and another species

While previous results are already based on an unseen test set of hormone-gene links performed within a sound 5-fold cross-validation framework, we also wanted to test how well our models would predict for fresh hormones that were never seen in the training/testing part of the cross-validation framework. In other words, we tested BioEmbedS on an independent dataset of hormone-gene pairs pertaining to “unseen” external hormones that were not used to train/evaluate the model (as these hormones had too few gene associations to be considered eligible in the final model building step and hence also every inner/outer CV iteration; see Algorithm 1 and associated text for details). This dataset contained 17 hormones forming 151 associated hormone-gene pairs and 148 non-associated hormone-gene pairs. Although, this dataset does not belong to the same distribution as the one used to train our model due to very few gene associations per hormone, BioEmbedS performed reasonably on this dataset obtaining accuracy of 65%, F1-score of 59%, and Area under ROC curve of 0.72 (Figure 3(b)). Similarly, we applied BioEmbedS-TS on a set of hormone-source/target gene pairs from 40 unseen hormones, forming 501 hormone-target gene pairs and 188 hormone-source gene pairs. It was able to correctly classify these pairs with 69% accuracy and area under PR curve of 0.43 (Figure 3(d)).

We also wanted to assess how well our model trained on one species (human) generalizes to make predictions in another species (mouse). That is, we applied BioEmbedS model trained on hormone-gene pairs from the HGv1.human dataset (referred to as HGv1 so far in the text) to assess how well it predicts hormone-gene associations in mouse (i.e., recovers the relations in HGv1.mouse dataset). Our model was able to achieve a reasonable accuracy of 71% and F1-score of 73%. Similary, when BioEmbedS-TS model trained on HGv1.human was used to classify hormone-associated genes in mouse into source and target genes, we were able to achieve a reasonable accuracy of 83%, and a F1-score of 78% for source genes and 87% for target genes. These results show that our BioEmbedS* models trained using human data generalize well to an organism other than human.

### 3.6 Novel gene predictions are enriched for the corresponding hormone-related diseases

The promising performance of BioEmbedS seen so far encouraged us to apply BioEmbedS to predict association between each hormone in HGv1 and all 19,318 human protein-coding gene symbols [3]. We were able to predict many novel hormone-gene links not captured in HGv1, comprising new links to any of the 1,453 genes in HGv1 (Table 2) or the remaining “out-of-HGv1” genes which were never seen during training/testing of our model. To validate these predictions, we tested whether the set of predicted genes for a hormone (at a default SVM probability score cutoff; see Methods) was enriched for diseases already known to be related to the hormone. We found this was indeed the case for 16 of the 34 primary hormones (Suppl Dataset D1a, with primary indicating hormones in HGv1 considered eligible for training the final model as already defined in Methods), and 9 of the 17 unseen external hormones (Suppl Dataset D1b, with unseen referring to the remaining hormones in HGv1 with very few gene associations). These results pertaining to hormones affecting a subset of tissues is shown in Figure 4. For instance, all the insulin predicted genes are indeed significantly enriched for “Diabetes Mellitus”, a disease term that is also recorded as insulin-related in the two hormone-disease “ground-truth” sources that we considered (Endocrine Society [https://www.hormone.org] and DisGeNET [22] resources; see Suppl Dataset D1a).

**Figure 4:**
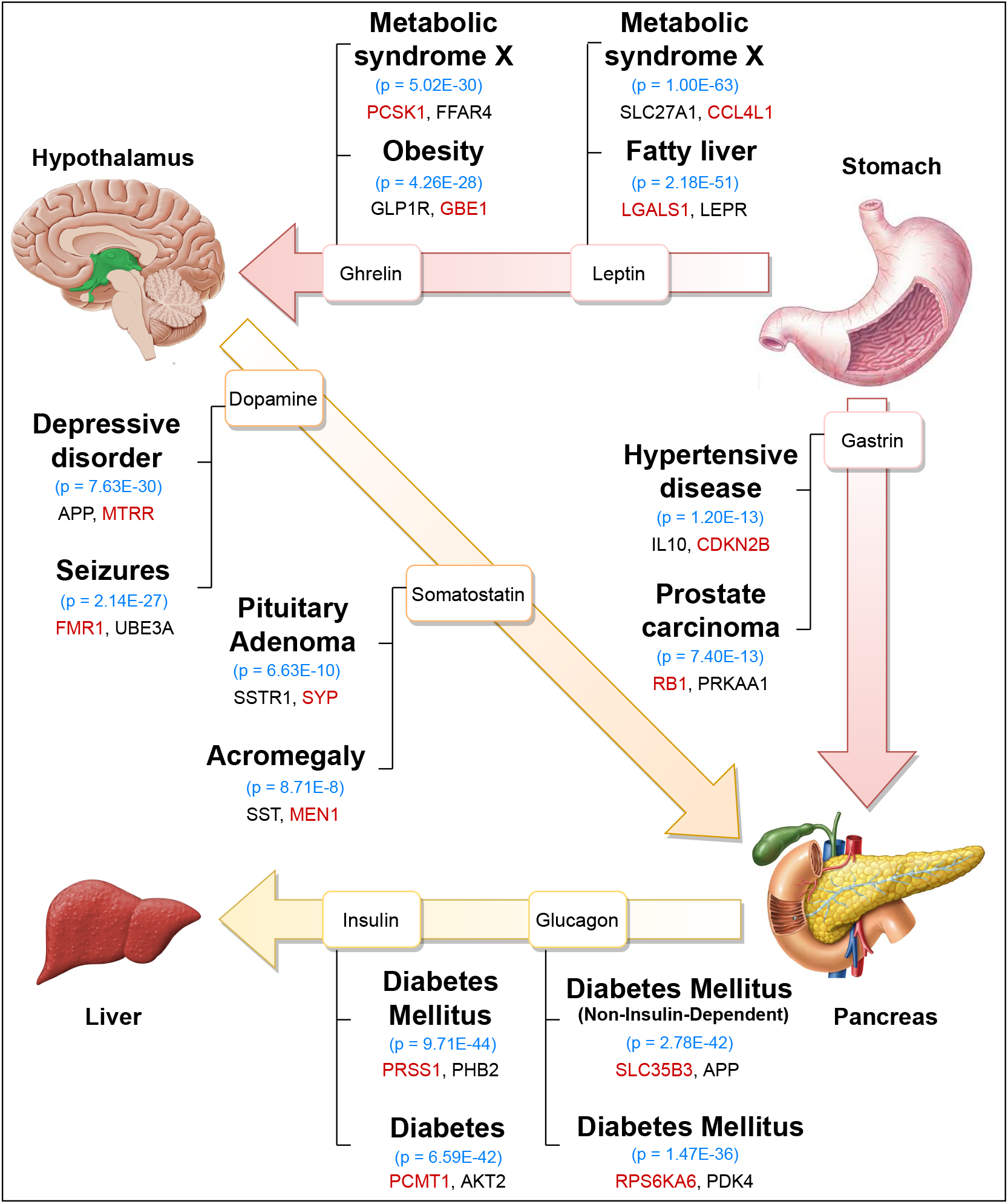
Inter-tissue communication: Example of a multi-tissue system with inter-tissue edges indicating hormonal signaling. BioEmbedS predictions for different hormones are enriched for the indicated diseases (top two are shown, along with disease enrichment P-values). Shown alongside each tissue-tissue link are examples of known disease genes that are also genes we predicted for a hormone (with black marking the genes in HGv1 dataset, and red the novel out-of-HGv1 genes). Tissue/organ pictures’ sources are given in Acknowledgments.

For well-studied hormones such as insulin, we repeated the above analysis on only the novel predictions (i.e., predicted hormone-gene links not in HGv1) to test if the disease term enrichments were driven not just by known hormone-specific genes in HGv1 but also by the novel predicted genes. For insulin, BioEmbedS predicted genes overlapped with 691 of the 1,507 Diabetes Mellitus genes recorded in DiSGeNET (disease enrichment *P* = 9.55 × 10^−40^), and 534 of these 691 overlapping disease genes were novel predictions (disease enrichment *P* = 4.85 × 10^−9^). This trend of enrichment of predicted novel genes for the corresponding diseases can also be seen visually in Figure 5 across a range of cut-offs applied on the SVM score to call predictions – specifically, the curve for novel gene predictions is better than that of a random classifier for insulin and other hormones, and follows closely the overall gene-curve for the most part. Furthermore, a more stringent subset of novel BioEmbedS predictions involving totally unseen out-of-HGv1 genes also got validated by a similar disease enrichment analysis

**Figure 5:**
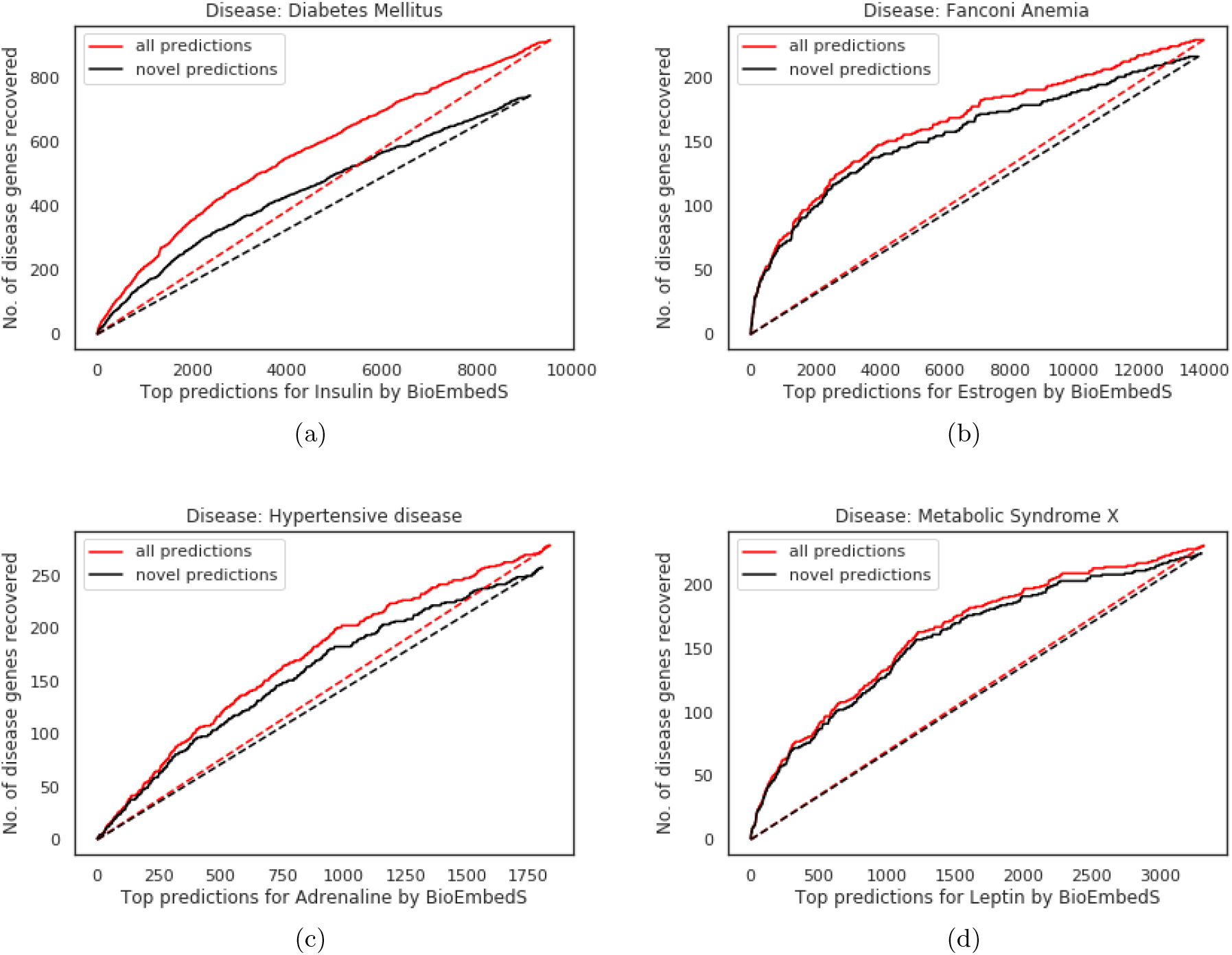
Disease enrichment in novel gene predictions: Curves showing the number (no.) of known disease genes (y-axis) recovered in top-*k* predicted genes (as per SVM score ranking; x-axis) of the corresponding hormone; focusing on all (red) vs. novel (black) predicted genes of the hormone. Our model (solid curves) performs better than chance recovery of disease genes by a random classifier (dashed lines). Only genes predicted for a hormone with SVM score > 0 are considered here; all protein-coding genes are considered in Suppl Figure S2.

## 4 Discussion

This work elucidates the computational problems and challenges in the emerging area of inferring cross-tissue signaling interactions from biomedical literature, and presents a first approach BioEmbedS to specifically predict hormone-gene associations from biomedical literature with reasonably good absolute accuracy and also comparable or better performance than other popular alternatives such as STRING. Our BioEmbedS and BioEmbedS-TS models are enabled by a ground-truth dataset HGv1 that we carefully complied and balanced across different stratifications of the training data in the space of mapped word embeddings of hormone names and gene symbols. The better performance of our method over other alternatives follows from our models being the first systematically developed ones to make hormone-gene predictions, and other methods being literature mining based relation extraction methods that are generic or designed for other prediction tasks such as disease-gene prediction.

Our HGv1 dataset can be viewed as a two-layered/bipartite graph (with HGv1 hormone-gene relations being the edges/links between nodes in the hormone layer and the gene layer), hence our hormone-gene prediction task can be viewed as a link prediction problem in bipartite graphs [17]. Current link prediction methods for bipartite or more general graphs [19, 16], such as ones based on pairwise node similarity, utilize attribute data available at the nodes and/or the structural connectivity/neighborhood of nodes based on existing links in the graph to predict new links. We decided to utilize only the attribute data (word embeddings) of nodes to do hormone-gene bipartite link prediction, since our HGv1 graph doesn’t have rich structural connectivity. In detail, a large number of genes in our HGv1 dataset (1,102 of the 1,453) are uni-hormone genes connected to only one hormone, and removing the few other multi-hormone genes disconnects the HGv1 bipartite graph into several connected components (one per hormone).

A caveat in this work worth mentioning is that randomly selected genes for a hormone need not be truly negative examples. Our model may also predict false associations based on high co-occurrence or context word similarity but no functional relationship, and a careful set of negative examples can help mitigate this issue. Our balancing strategy using SMOTE is also not without its pitfalls, especially when applied to high-dimensional data [13], and we coupled it with undersampling to carefully balance our training data across different hormones that are represented in the literature to different extents and to address imbalance in the number of source vs. target genes. Nevertheless, good accuracy of our model on different unseen test sets and on an organism other than human, and our disease enrichment analysis of novel gene predictions taken together suggest that our model predictions are indeed generalizable to make novel predictions about inter-tissue gene signaling mediated by a hormone.

Future work would focus on integration of novel hormone-gene predictions with independent data such as multi-tissue genomic data [10], and systematic interpretation/validation of non-coding gene predictions for hormones (such as the preliminary lncRNA (long non-coding RNA) predictions that we provide in our website). Future work could also try integrating literature information with data on protein-protein, protein-DNA or other interactions among genes and gene products to further improve model performance. A special feature of our ground-truth hormone-gene dataset HGv1 is its applicability to organisms beyond human and mouse as it is based on species-agnostic Gene Ontology terms, and this bodes well for extending our work to other multicellular organisms in the future. We hope this work stimulates more work along these lines on cross-tissue signaling and advances the field forward to developing whole-body, cross-tissue gene networks for different organisms.

## Acknowledgments

We thank Arjun Sarathi for help in assembling the HGv1 dataset, and BIRDS (Bioinformatics and Integrated Data Science) group members for their valuable suggestions and reviews for this work, and Sanga Mitra and Sugyani Mahapatra in particular for their careful reviews, and Philge Philip for help with the code repository. We thank Balaraman Ravindran and Harish Guruprasad from IIT Madras, and Praveen Anand from nference for their valuable inputs on earlier stages of the project. The research presented in this work was supported by WT/DBT grant IA/I/17/2/503323 awarded to MN.

Pictures of tissues/organs in Figure 4 are taken from the following sources - Liver: https://www.medicalnewstoday.com/articles/305075, Pancreas: https://www.shutterstock.com/image-illustration/anatomy-drawing-showing-pancreas-duodenum-gallbladder-1396704593, Hypothalamus: https://quizlet.com/289736618/human-brain-midsection-diagram/, Stomach: https://zen.yandex.ru/media/wowfacts/neskolko-faktov-o-jeludke-5b916e7cc586d600aa836cc5.

## Author Contributions

AJ and MN formulated the study and overall modeling approaches; AJ implemented the training/testing of all primary models and performed associated analyses; TK and MN compiled the HGv1.human dataset, and provided inputs on the modeling approaches; TK assembled the HGv1.mouse dataset, and studied novel gene predictions at the hormone and inter-tissue level; MR performed associated analyses of secondary modeling approaches, visualized embeddings, and developed the website with inputs from AJ; TL performed disease and pathway enrichment, hierarchical clustering, and exploratory lncRNA analyses; AJ, MN, TK, TL and MR interpreted results; AJ, MN and TK wrote the manuscript; MN guided and supervised the study.

## Supplementary Information

### 1 Supplementary Methods

#### 1.1 Performance Metrics

As explained in the main text, the positive and negative class in our binary classification problems refer respectively to: association and non-association of a hormone-gene pair in the BioEmbedS setting, and hormone-source gene pair and hormone-target gene pair association in the BioEmbedS-TS setting. Applying standard definitions to these settings yields the following definitions, with number abbreviated as “#”.

True Positives (TP): # of positive hormone-gene pairs predicted as positive by BioEmbedS; # of hormone-source gene pairs predicted correctly by BioEmbedS-TS.
True Negatives (TN): # of negative hormone-gene pairs predicted as negative by BioEmbedS; # of hormone-target gene pairs predicted correctly by BioEmbedS-TS.
False Positives (FP): # of negative hormone-gene pairs predicted as positive by BioEmbedS; # of hormone-target gene pairs predicted as hormone-source gene pairs by BioEmbedS-TS.
False Negatives (FN): # of positive hormone-gene pairs predicted as negative by BioEmbedS; # of hormone-source gene pairs predicted as hormone-target gene pairs by BioEmbedS-TS.

We evaluate our classifiers on the following performance metrics derived from the above counts.

1. Precision: In the context of BioEmbedS classifier, it represents the proportion of predicted hormone-gene pairs (TP + FP) that are actually correct as per the HGv1 dataset (TP). In the context of BioEmbedS-TS, it indicates the proportion of predicted source genes that are truly the source genes as per the HGv1 dataset.

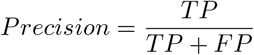
2. Recall: In the context of BioEmbedS, it is the ratio of hormone-gene associations that our model can predict (TP) to the total associations present in the HGv1 dataset (TP + FN). In the context of BioEmbedS-TS, it is the ratio of source genes that our model recovers to all source genes present in the HGv1 dataset.

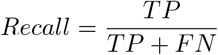
3. F1-score: It is the harmonic mean of Precision and Recall scores.
4. Accuracy: It indicates out of all the model’s predictions, how many are correct predictions.

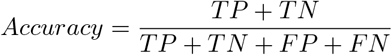
5. Cohen’s Kappa score: It indicates how often the model’s predictions and the actual HGv1 labels for all considered hormone-gene pairs agree relative to random chance agreement, and is a useful metric for classification with imbalanced datasets [2].
6. ROC-AUC: The area under the *Receiver Operating Characteristics* (ROC) curve, which plots TPR (true positive rate or recall 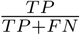) on the y-axis against FPR (false positive rate or 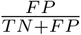) on the x-axis, with different points on the curve based on different cutoffs applied on the model scores to make positive vs. negative class predictions [1].
7. PR-AUC: The area under the *Precision-Recall* (PR) curve, which plots precision on the y-axis against recall on the x-axis, with different points on the curve again based on different cutoffs applied on the model scores to make positive vs. negative class predictions [1].

#### 1.2 Hyperparameters of BioEmbedS classifiers

Besides choosing SVM (Support Vector Machines) and RF (Random Forests) as our primary classifiers for use with the BioEmbedS model (see main text and Suppl Table S1), we also tried other secondary choices of classifiers (see Suppl Table S2). Hyperparameters of these primary and secondary classifiers are given below, and are implemented using the *Scikit-learn* machine learning framework in Python [3]. There were no hyperparameters to choose for simpler models like logistic regression.

SVM: The range of hyperparameter values considered for the SVM classifier are as follows. For kernel functions, we tried RBF (Radial Basis function) and polynomial kernel types. The model complexity parameter C had 9 equally spaced values between −4 to 4 in the log space. Gamma parameter for RBF kernel had 12 equally spaced values between −9 to 2 in the log space. Degree parameter for the polynomial kernel had values 2, 3, 5 and 7. In each fold, polynomial kernel with degree = 3 and *C* = 1, was chosen as the best classifier based on scores on the validation set. We also choose this hyperparameter setting of SVM as our final classifier model to make novel predictions.
RF: The range of hyperparameter settings considered for the sklearn implementation of the RF classifier are as follows. For the number of trees in the forest, we tried 7 arbitrarily pre-selected values from 100 to 1600; and for the maximum depth of each tree, we tried 9 pre-selected values from 120 to 360. We let the minimum number of samples required to be at a leaf node to be 1, 2, or 4; and the minimum number of samples required to split an internal node to be 2, 3, 5, or 7.
Neural Networks: The Neural Networks have 2 hidden layers. The number of units in each layer was sampled among 32, 64 and 128 units. Four values of learning rate were tried out between 0.0001 and 0.1, and the regularisation parameter was chosen among 10^*−*3^, 10^*−*4^, and 10^*−*5^. From all these possible configurations, the combination of parameters that gave the best results were chosen.
XGBoost: For the XGBoost model, 5 values of learning rate were sampled between 0.03 and 0.3 in the log-space, 5 values of maximum depth were sampled between 2 and 6, and 5 values of the number of estimators were sampled between 100 and 150 in the linear space. From these, the combination of parameters that gave the best results were chosen.
Decision Trees: Five values of maximum depth were sampled between 2 and 6, and five values of the number of estimators were sampled between 100 and 150 in the linear space.

## 2 Supplementary Tables

**Table S1:** BioEmbedS performance across 5 test folds: Results are using the 5 test folds of our cross validation framework using the best primary classifier (which turned out to be a SVM classifier with degree-3 polynomial kernel and *C* = 1 as mentioned in Suppl Methods 1.2).

| Fold | Precision | Recall | F1-score | Accuracy | ROC-AUC | PR-AUC | Kappa Score |
| --- | --- | --- | --- | --- | --- | --- | --- |
| 1 | 0.70 | 0.76 | 0.73 | 0.71 | 0.78 | 0.76 | 0.43 |
| 2 | 0.70 | 0.71 | 0.70 | 0.70 | 0.76 | 0.76 | 0.40 |
| 3 | 0.67 | 0.70 | 0.69 | 0.68 | 0.74 | 0.71 | 0.36 |
| 4 | 0.71 | 0.77 | 0.74 | 0.73 | 0.79 | 0.76 | 0.46 |
| 5 | 0.70 | 0.72 | 0.71 | 0.70 | 0.76 | 0.74 | 0.40 |

**Table S2:**
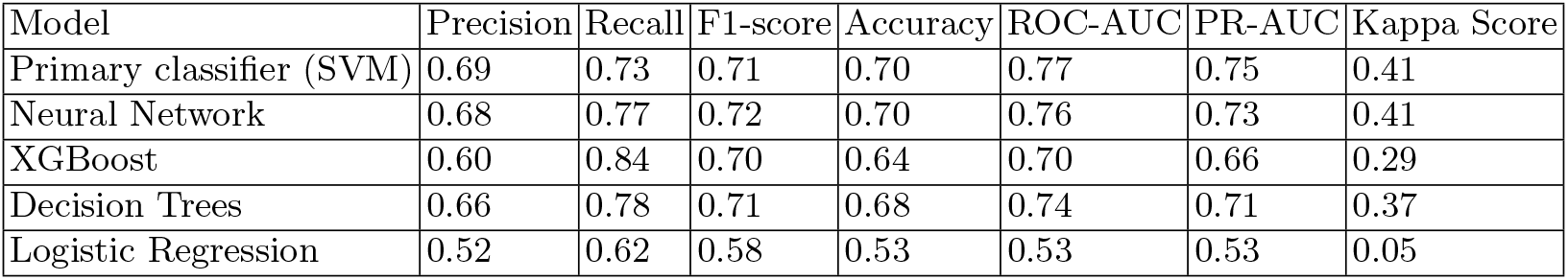
BioEmbedS performance for different choices of classifiers: Performance reported is average across the 5 cross-validation test folds – for instance SVM’s performance is average of performance reported in Table S1. It is evident that the SVM classifier achieved better or comparable performance relative to other classifiers. It is also clear that simpler models like Logistic Regression could not capture patterns in the dataset, and higher order function approximators like Neural Networks or algorithms like SVM provide better results.

## 3 Supplementary Figures

**Fig. S1.**
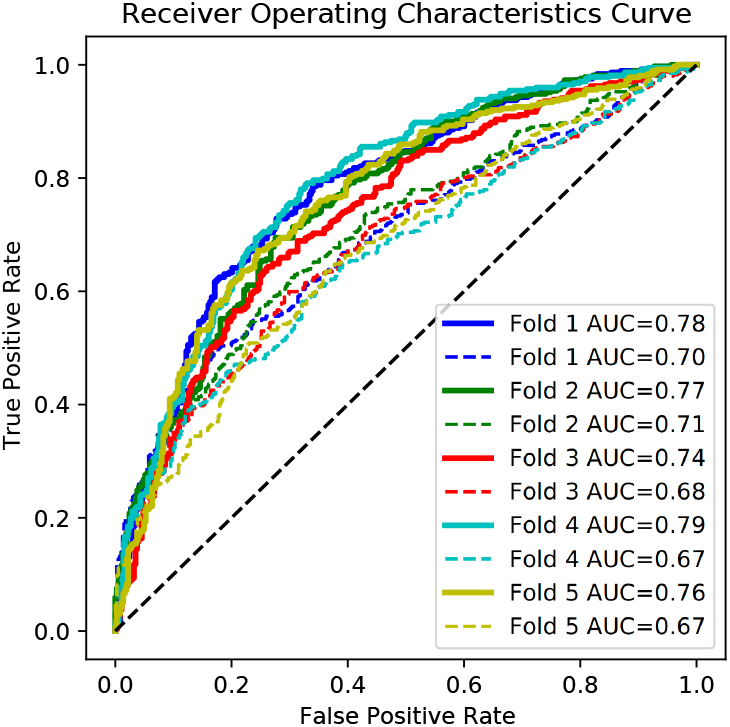
Cosine similarity and BioEmbedS performance: ROC curves for hormone-gene predictions using unsupervised cosine similarity based method (dashed lines), and our supervised method BioEmbedS based on the SVM classifier (solid lines).

**Fig. S2.**
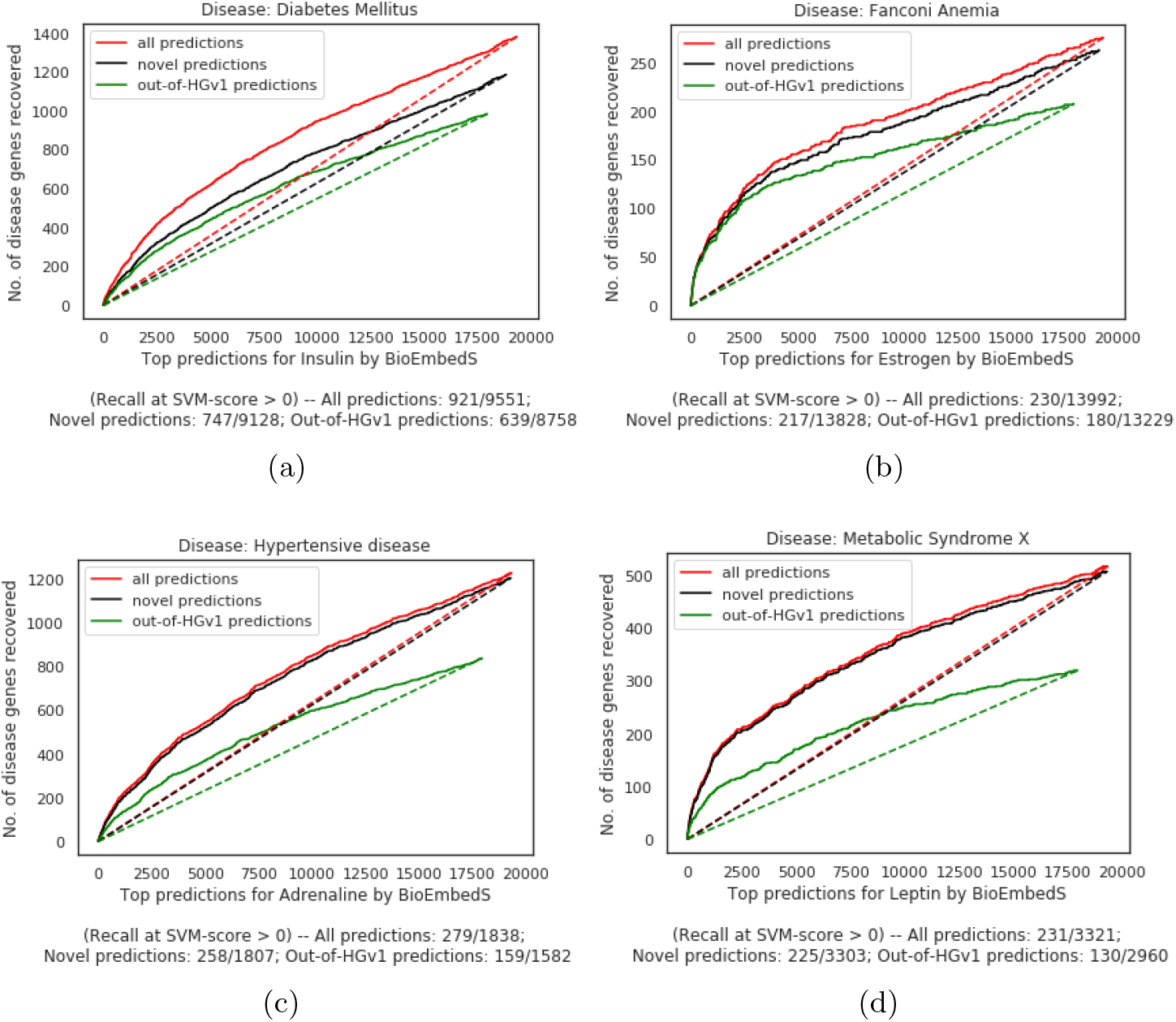
Disease enrichment in novel and out-of-HGv1 gene predictions: Curves showing the number (no.) of known disease genes (y-axis) recovered in top-*k* predicted genes (as per SVM score ranking; x-axis) of the corresponding hormone; focusing on all (red) vs. novel (black) vs. out-of-HGv1 (green) predicted genes of the hormone. To clarify these terms for a given hormone, ‘all” refers to predictions made for all 19,318 protein-coding genes; “novel” refers to a subset of all predicted genes such that the hormone-gene **pair** is not in HGv1; and “out-of-HGv1” refers to a subset of the novel predicted genes such that the **gene** is also not in HGv1 (i.e., the predicted gene is not associated with any hormone in HGv1, and hence totally absent from HGv1 and unseen during model building). Our model (solid curves) performs better than chance recovery of disease genes by a random classifier (dashed lines). Below each hormone’s disease enrichment plot, the number of disease genes overlapping the predicted genes at SVM score > 0 for the hormone is shown as a ratio (# overlapping disease genes / # predicted genes). It is evident that even after removing the hormone-associated HGv1 genes, a significant number of disease-related genes are left and they get predicted towards the top by BioEmbedS (black curves). Moreover, BioEmbedS performs well on the totally unseen out-of-HGv1 genes too (green curves).

## 4 Supplementary Datasets

Supplementary datasets are available at this link.

Suppl Dataset D1a: Disease enrichment analysis of predicted genes for the (34) primary hormones in the HGv1 dataset.

Suppl Dataset D1b: Disease enrichment analysis of predicted genes for the (17) unseen/external hormones in the HGv1 dataset.

## Footnotes

1 https://www.hormone.org/your-health-and-hormones/glands-and-hormones-a-to-z, accessed Jul 23, 2019.

2 Human-mouse homology mapping done via MGI Batch Query (http://www.informatics.jax.org/batch), accessed Nov 11, 2020.

3 BioWordVec model/embeddings are downloaded from https://github.com/ncbi-nlp/BioSentVec.

4 https://github.com/BIRDSgroup/BioEmbedS

